# The optimization of mRNA expression level by its intrinsic properties—insights from codon usage pattern and structural stability of mRNA

**DOI:** 10.1101/202200

**Authors:** Manish P Victor, Debarun Acharya, Tina Begum, Tapash C Ghosh

**Author notes:** Authors for correspondence: Debarun Acharya, Bioinformatics Centre, Bose Institute, Kolkata, West Bengal, India, Tapash C Ghosh, Bioinformatics Centre, Bose Institute, Kolkata, West Bengal, India.

## Abstract

The deviation from the uniform usage of synonymous codons is termed codon usage bias. A lot has been explained from the translational viewpoint for the observed phenomenon. To understand codon usage bias from the transcriptional perspective, we present here a holistic picture of this phenomenon in *Saccharomyces cerevisiae*, using both wild type and computationally mutated mRNAs. Although in wild type, both codon usage bias and mRNA stability positively regulate the gene (mRNA) expression level and are positively correlated with each other, any deviation from natural situation breaks such equilibrium. Computational examination of mRNA sequences with different sets of synonymous codon composition reveals that in mutated condition, the mRNA expression becomes reduced. Furthermore, constraining codon usage pattern to wild type and carrying out randomization of codons resulted in less stable mRNA. Further, we realized a Boolean Expression explaining the gene expression under various conditions of bias and mRNA stability. These studies suggest that selection of codons is favored for regulation of gene expression through potential formation of messenger RNA structures which contribute to folding stability. The naturally occurring codon composition is responsible for optimization of gene expression, and under such composition, the mRNA structure having highest stability is selected by nature.

## 1. Introduction

Codons are the combination of three DNA or RNA nucleotides that encodes an amino acid and supplies it to the polypeptide chain or stops protein synthesis during translation. The codons sum to a total of 61 in number that encodes 20 amino acids. With Tryptophan and Methionine as exceptions, all other amino acids are coded by two to six codons, even in the same polypeptide or protein. All codons responsible for coding the same amino acid are referred to as synonymous codons^1^. In order to understand these synonymous codons let us consider the antioxidant tri-peptide glutathione, made up of cysteine, glutamic acid, and glycine which can be encoded by sixteen different coding sequences. Even with a lot of coding sequences possible for every protein theoretically, only one of them is selected by nature^2^. Such deviation from uniform usage of the codons is known as codon usage bias (CUB) that refers to the differences in the frequency of occurrence of synonymous codons in coding DNA^1^. It incites queries such as (a) Does codon bias facilitate accuracy in gene expression? As polypeptide/proteins that have important functional aspects in an organism. So, erroneous polypeptides/proteins will be harmful to organisms. Repeated use of a certain set of codons may reduce the chances of misincorporation of amino acids in the polypeptide/protein. However, it raises yet another query— (b) under the high demand of protein do such bias help in high gene expression level? The underlying thought is that under certain conditions, the cellular demand for a protein may increase. Thus, translation needs to escalate with the demand. A small bunch of tRNAs may reduce the screening procedure and facilitate rapid expression with fidelity. Because transcription and translation are articulated processes, it raises another question, i.e. (c) Is codon bias, helpful in any way at the transcript level, as it is the connecting link between protein and DNA? In the pursuit of inferring these queries, mutation-selection-drift balance model has been put forward, which highlights the influence of selection and mutation in determining the codon usage pattern in both highly and lowly expressed genes^1,3–5^. For lowly expressed genes, there is no definitive evidence that they have a preference for rare codons, which has been well accounted by the high synonymous substitution rate for weakly expressed genes in Enterobacteria under relaxed selection pressure^6^. So, for the codons, at one end neutral pressure propounds the unvarying effect throughout the genome for its usage. Under neutral pressure, mutational bias comes into picture which is known to differ between organisms. Hence, this leads to the variation in the codon bias across organisms^1^. It is to the non-randomness in mutational pattern that illustrates for the prevailing codon usage bias with GC-content variations across genomes^1,7,8^.

On the other end from the selectionist perspective, it implies that since highly expressed genes are used by the organisms more often than lowly expressed genes, it is more likely for the former to show greater codon usage bias. Studies of codon usage pattern in Enterobacteria, *Escherichia coli* and Yeast show the predominant use of optimal codons in highly expressed genes, compared to the uniform use of all synonymous codons in weakly expressed genes^5,9^. It has also been shown by the translational selection hypothesis that codon usage is very closely entwined with the tRNA pool, preventing translation machinery from incorporating incorrect amino acid(s)^10,11^. These elucidations have been restrained only at the translational level, i.e., when demand is high bias helps in high gene expression level with fidelity.

In the bigger picture, the transcriptional domain of Central Dogma seems to be margined out. Ribonucleic acid has played an essential role since the inception of life, as is the bridge between DNA and protein. In a recent study by Mao *et al.*, mRNA secondary structures were considered to facilitate the appropriate abundance and binding of ribosomes^12^. In addition to fidelity and the rate of translation facilitated by codon usage bias, it may also be the determinant of secondary structure. mRNA stability refers to the maintenance of mRNA structural integrity. mRNA stability arising out of its innate secondary structures play a major role in influencing protein abundance^13^, transcript splicing^14^, is responsible for halts in protein synthesis^15,16^, determinants of protein secondary structure^17–20^, and also play a role in regulating gene expression level as well as the accuracy in expression^12,21,22^. Studies have also shown preference of codons pairing to high-abundance tRNAs in the regions of high mRNA secondary structure content, while codons read by significantly less abundant tRNAs are situated in lowly structured regions of the mRNA^23^. As these suggest, both translation and transcription are not exclusive processes and modulated by mRNA secondary structure; it has paved the way for studies of mRNA stability on codon bias determination. To address the third question of CUB in influencing transcript levels, Edoardo Trotta presented the influence of transcriptional forces on codon usage bias that improves gene expression by optimising mRNA folding structure^2^.

In this study, we tried to address how CUB and mRNA structural stability is associated with its expression. The results astonishingly indicate a harmonious articulation of trio, i.e., codon composition, mRNA secondary structure stability, and mRNA abundance. We found that even under the high stability and narrow selection of codon pool does not suffice for the appropriate selection for gene expression level. Our study here confirms that the existent codon composition has not a mere influence of translational domain only, as transcription and translation are both articulated process. Rather, it indicates that both mRNA stability and meticulous codon usage fine tunes the apt gene expression level.

## 2. Materials and Methods

### 2.1. Data collection and filtering

The exonic sequences of 5917 *Saccharomyces cerevisiae* ORFs were downloaded from Saccharomyces Genome Database (http://downloads.yeastgenome.org/sequence/S288C_reference/orf_dna/orf_coding.fasta.gz). The erroneous sequences were checked with CodonW (version 1.4.4, https://sourceforge.net/projects/codonw/)^24^ and removed. Finally, we obtained 5860 ORFs as our dataset.

### 2.2. mRNA and Protein abundance data

The number of mRNAs molecules per cell were collected from genome wide expression analysis for 6180 genes by Holstege *et al.* (http://web.wi.mit.edu/young/pub/data/orf_transcriptome.txt)^25^. We also used a more up-to-date large scale expression data set for validation purpose from supporting information Table S1 of Pelechano *et al.* ^26^ From this table, we extracted the transcripts per cell values by Nagalakshmi *et al.* and Miura *et al.* and obtained the gene expression values for 5086 genes^27,28^. We extracted the protein abundance dataset comprising 5701 genes from Arava *et al* ^29^.

### 2.3. mRNA stability calculation

The mRNA stability was calculated using two separate approaches. Firstly, we obtained the experimentally determined mRNA secondary structures of the Baker’s Yeast *Saccharomyces cerevisiae* from Kertesz *et al.*,^30^. Similar to their analysis, we used the “Nucleotide resolution raw reads” scores (http://genie.weizmann.ac.il/pubs/PARS10/pars10_catalogs.html) representing the nucleotides being in paired (stated as V1 reads) and unpaired (stated as S1 reads) states to calculate mRNA folding score by using this formula-

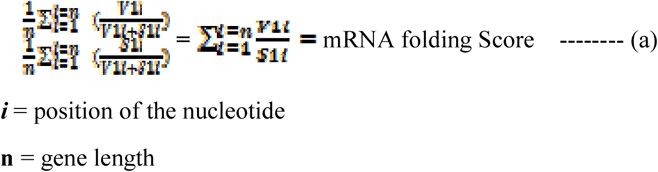

As the efficiency of enzymes varies, the ratios suggested by Kertesz *et al.*^30^ for V1 and S1 reads are relative and are meaningful only when compared among nucleotide sites^31^. Thus, we have calculated a mRNA folding score for each gene where numerator and denominator are the arithmetic mean of the probabilities of nucleotides being in the paired and unpaired conditions. As the numerator represents the paired state so, higher the folding score, greater the fraction of paired sites in the mRNA, consequently stronger is the overall folding of the mRNA. The folding score ranges from 0.4 to 2.4 in our dataset.

The second approach to predict the folding free energies of the sequences we used the RNAfold attribute of the Vienna RNA package Version 2.0 ^32^. We used the folding temperature of 30 ◻C as the previous experiment was carried under this temperature. The prediction is based on a thermodynamic model with the assumption that the secondary structure of an mRNA is determined by the minimum free energy ^33^. As long range interactions are scarce ^34,35^ and computationally demanding ^36^. We estimated the local folding energies of the ORFS by taking the sliding windows of 40 nucleotides with a step size of 1 nucleotide ^13^. Specifically, mean predicted local folding energies reported in this paper are calculated for the entire transcript of each gene.

So,

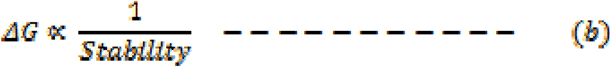

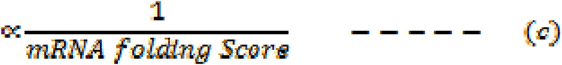

### 2.4. Generating Mutated Genome

Two types of computationally generated mutated genome of *Saccharomyces cerevisiae* were generated by using two in-house PERL scripts. Firstly, we incorporated different codon composition with fixed codon usage from 20 codons for all the 20 amino-acids to all 61 codons for 20 amino-acids (i.e. 20, 30, 40, 50 & 61) with different levels of nucleotide contents (AT-rich, when at least two of the three nucleotides are either A or T; GC-rich, when at least two of the three nucleotides are either G or C). We also made subgroups of different AT-GC composition for each of the groups, where applicable. Thus, under every group for each mRNA, four different values were obtained for the same parameter.

Secondly, by using another PERL script, we randomly shuffled the synonymous codons within the mRNA, keeping the ENC same as the native form. For each mRNA, we made one hundred such random synonymous codon substituted combinations and calculated the mRNA stability. To derive mRNA folding score here, we implemented the similar method as above. For each window size of 40bp used in calculating the folding free energy, every base was monitored in it of being in a paired and unpaired state. We considered 0.65 as the cut-off value of pairing probabilities for each nucleotide. Nucleotide bases having values greater than or equal to 0.65 was considered as ‘paired’ and below it was included in the unpaired state. We have also used a more stringent cut-off value (0.80) for the same analysis.

### 2.5. Softwares

CodonW (version 1.4.4) (http://codon.sourceforge.net/) was used to calculate codon bias indices: Codon Adaptation Index (CAI) and the Effective Number of Codons (Nc). The statistical analysis was carried using IBM SPSS statistics 22 using parametric and non-parametric tests. All the correlations reported in this study are Spearmen correlation. As dependence of only ENC is to be found on other factors, linear regression analysis was carried out. In all statistical analysis, we used 95% level of confidence as a measure of significance.

## 3. Results and Discussion

### 3.1. The relationship between mRNA stability and mRNA expression level

The stability of messenger RNA throughout this study refers to the structural integrity of this biomolecule and not to be confused with its half-life or degradation rate^12,13,16,21,37^. We calculated the stability of mRNAs using two approaches. Firstly, as a mRNA is a stretch of ribonucleotides with many secondary structures like loops, stems, hairpins, etc., endowed with the Watson-Crick base pairing, such structural integrity is represented as mRNA folding score and is used as a measurement of mRNA stability^2,30^. Secondly, at the molecular level, the stability of any molecule can be calculated using its free energy (ΔG)^37^, where a lower ΔG represents higher stability of that molecule. Hence, for a more precise measurement, the mRNA folding free energy was also used independently to assess the structural stability of the *Saccharomyces cerevisiae* mRNAs. We obtained the mRNA expression level (mRNA abundance) independently from the two datasets of Holstege *et al.^25^* and Pelechano *et al.^26^.* To understand the interplay among mRNA stability and mRNA expression level we correlated the two factors. There exists a significant positive correlation between mRNA folding score and mRNA expression level for all the datasets *ρ*=0.394; *P*=3.049×10^−107^; *n*=2886) and *ρ*=0.346; *P*=2.202×10^−74^; *n*=2606) (Table 1, Table2). Using folding free energy to measure the mRNA stability, it exhibited a significant negative correlation with expression level for all datasets (*ρ*=−0.342; *P*=3.545×10^−141^; n=5139) and (*ρ*=−0.277; *P*=2.689×10^−86^; *n*=4841) (Table 1, Table2). Both results indicate that the more stable mRNA becomes, its expression value increases. Additionally, as the stability of an mRNA is dependent on its structural features, i.e., the ribonucleotides sequences, any change to this sequence is likely to alter the structure and the stability. As we know that there are many different codon composition that encodes for the same amino acids (synonymous codons), the structural stability of an mRNA may be affected by differential use of codon, without any alteration to its encoded protein sequence. Therefore, for mRNAs that encode for same protein but are structurally different the mRNA expression levels may differ. Now, as we know that the preferential use synonymous codon is described as codon usage bias, such bias may cause alteration in mRNA stability as well as its expression level.

**Table 1.** Correlation of mRNA stability with other parameters among Yeast genes. (Holstege *et al.* dataset ^25^)

| Parameters | Correlation Coefficient | *P*-value | N |
| --- | --- | --- | --- |
| Expression level | -0.342 | $3.545 \times 10^{-141}$ | 5139 |
| ENC | 0.108 | $1.405 \times 10^{-16}$ | 5859 |
| CAI | -0.160 | $4.788 \times 10^{-35}$ | 5859 |

6 1B. Rank correlation mRNA stability (folding score) with various parameters
| Parameters | Correlation Coefficient | *P*-value | N |
| --- | --- | --- | --- |
| Expression level | 0.394 | $3.049 \times 10^{-107}$ | 2866 |
| ENC | -0.329 | $1.574 \times 10^{-76}$ | 2994 |
| CAI | 0.410 | $5.808 \times 10^{-122}$ | 2994 |

**Table 2.** Correlation of mRNA stability with other parameters among Yeast genes. (Vincent Pelechano *et al.* dataset ^26^).

| Parameters | Correlation Coefficient | *P*-value | N |
| --- | --- | --- | --- |
| Expression level | -0.277 | $2.689 \times 10^{-86}$ | 4841 |
| ENC | 0.108 | $1.405 \times 10^{-16}$ | 5859 |
| CAI | -0.160 | $4.788 \times 10^{-35}$ | 5859 |

5 2B. Rank correlation mRNA stability (folding Score) with various parameters
| Parameters | Correlation Coefficient | *P*-value | N |
| --- | --- | --- | --- |
| Expression level | 0.346 | $2.202 \times 10^{-74}$ | 2606 |
| ENC | -0.329 | $1.574 \times 10^{-76}$ | 2994 |
| CAI | 0.410 | $5.808 \times 10^{-122}$ | 2994 |

### 3.2. Relationship between mRNA stability and codon usage bias

Studies done earlier have shown that genes with higher expression levels show biased usage of codons^12,38^. So, we tried to find any relationship between codon usage bias and mRNA stability, as mRNA stability and its expression level hold a significant positive correlation. We used the effective number of codons to calculate the codon usage bias. Effective number of codons (ENC or *Nc)* is the measurement of the state of codon usage biases (CUB) in genes and genomes and ranges from 20 (extreme bias using only one codon per amino acid) to 61 (no bias towards any codon)^39^. Although the determination of ENC values are uncomplicated, this measure is till date one of the best methods to show codon usage bias^40^. With folding free energy, ENC exhibits a notable positive correlation (*ρ*=0.108; *P*=1.405=10^−16^; *n*=5859). Similarly, with the folding score, ENC exhibits a significant negative correlation (*ρ*= −0.329; *P*=1.574×10^−76^; *n*=2994), both indicating an increase in mRNA stability for genes having higher CUB. Therefore, a biased usage of codons may lead to mRNAs with higher structural stability that encodes for the same polypeptide chain.

### 3.3. Manipulation of codon composition and its effect on mRNA expression level

From the previous analyses, it is quite clear that genes that express at high levels possess high levels of codon usage bias. To find out the influence of bias on the expression of all genes, we did the following study. We grouped the usage of codons under being AT-rich, GC-rich, and moderately AT-GC-rich. For each group, different numbers of codons were used ranging from 20(extremely biased) to 61 (nil biased). We hypothesize that if CUB is important for elevated gene expression, codon manipulation towards the more biased state should elevate the gene expression for all genes. However, the mRNA abundance under different sets of conditions seemed difficult to calculate. Thus, to arrive at appropriate values for mRNA abundance, we use the CAI values as a proxy. For all ranges of codons (17 categories) (Supplementary table S1) used we found that gene expression level (CAI) positively correlated with free folding energy and negatively with the folding score. This positively suggests that under different codon bias, mRNA abundance negatively changes with stability (Supplementary table 1). It means that with decreasing stability expression increases. However, the results from our dataset and earlier studies^12,13,31,37^ it seems quite impossible for the result to be true in natural situations. The possibility of such a condition seems unfavorable because to a minimum extent the stability of secondary structures lengthens its degradation procedure^41^. Without proper folding ribosomal binding will be affected and consequently, apt translation will be hampered^12^. Moreover, under folded structures, ribosomes stall, helping in the co-translational folding^20,42^ which will be highly compromised under reduced stability of the secondary structures of mRNA. In the mutated condition when higher or lower levels of CUB were applied to the mRNAs, the gene expression values (i.e., CAI) becomes decreased in either case (Table 3, Supplementary table S1). Here, we noticed that the average CAI for original sequences comes out to be 0.18 (Fig. 1). Whereas for all other cases in mutational analysis be it a highly stable set of genome or all of the codons consumed genome, the maximum average CAI amongst these categories comes to be 0.14 and minimum to be 0.07 (Fig.1). Therefore, it can be concluded that the existing codon usage pattern helps to achieve its maximum gene expression in yeast. Although biased usage of codons generally increases the gene expression, as highly expressed genes have high codon usage bias, this is not the case here. We found that any deviation from native codon usage may lead to reduced mRNA production (Table 3), which will lead to reduced production of proteins, as mRNA and protein expression have a strong positive correlation [(ρ=0.614; P=1.0×10-6; n=4833) and (ρ=0.612; P=1.0×10-6; n=4374)]. Therefore, it is quite clear that the when the system is forced to a more biased state, the expected increment in gene expression level does not happen, and hence, the native state has the highest selective advantage for maximising the gene expression level.

**Table 3.** This table shows the comparison between average CAIs from the wild type and the sequences generated computationally with different levels of codon compositions being used. All the other sequences show low CAI values as compared to the wild type sequences due to the usage of unconventional and rare codons. Here, numbers 20, 30 ….. Show the number of codons being used AT, GC, ATGC show the base
compositions of the genome. The ENC values for each class is also shown here.

| Codons/Genome | CAI<sub>AVG</sub> | ENC |
| --- | --- | --- |
| Native Sequence | 0.18 | 50.19 |
| 20AT | 0.07 | 20 |
| 20GC | 0.10 | 20 |
| 20ATGC1 | 0.08 | 20 |
| 20ATGC2 | 0.09 | 20.96 |
| 30AT | 0.13 | 26.51 |
| 30GC | 0.07 | 31.33 |
| 30ATGC1 | 0.08 | 33.34 |
| 30ATGC2 | 0.11 | 31.43 |
| 40AT | 0.14 | 41.6 |
| 40GC | 0.07 | 39.8 |
| 40ATGC1 | 0.11 | 41.47 |
| 40ATGC2 | 0.08 | 38.20 |
| 50AT | 0.12 | 53.28 |
| 50GC | 0.08 | 53.28 |
| 50ATGC1 | 0.09 | 54.25 |
| 50ATGC2 | 0.10 | 54.34 |
| 61 | 0.09 | 61 |

**Fig. 1.**
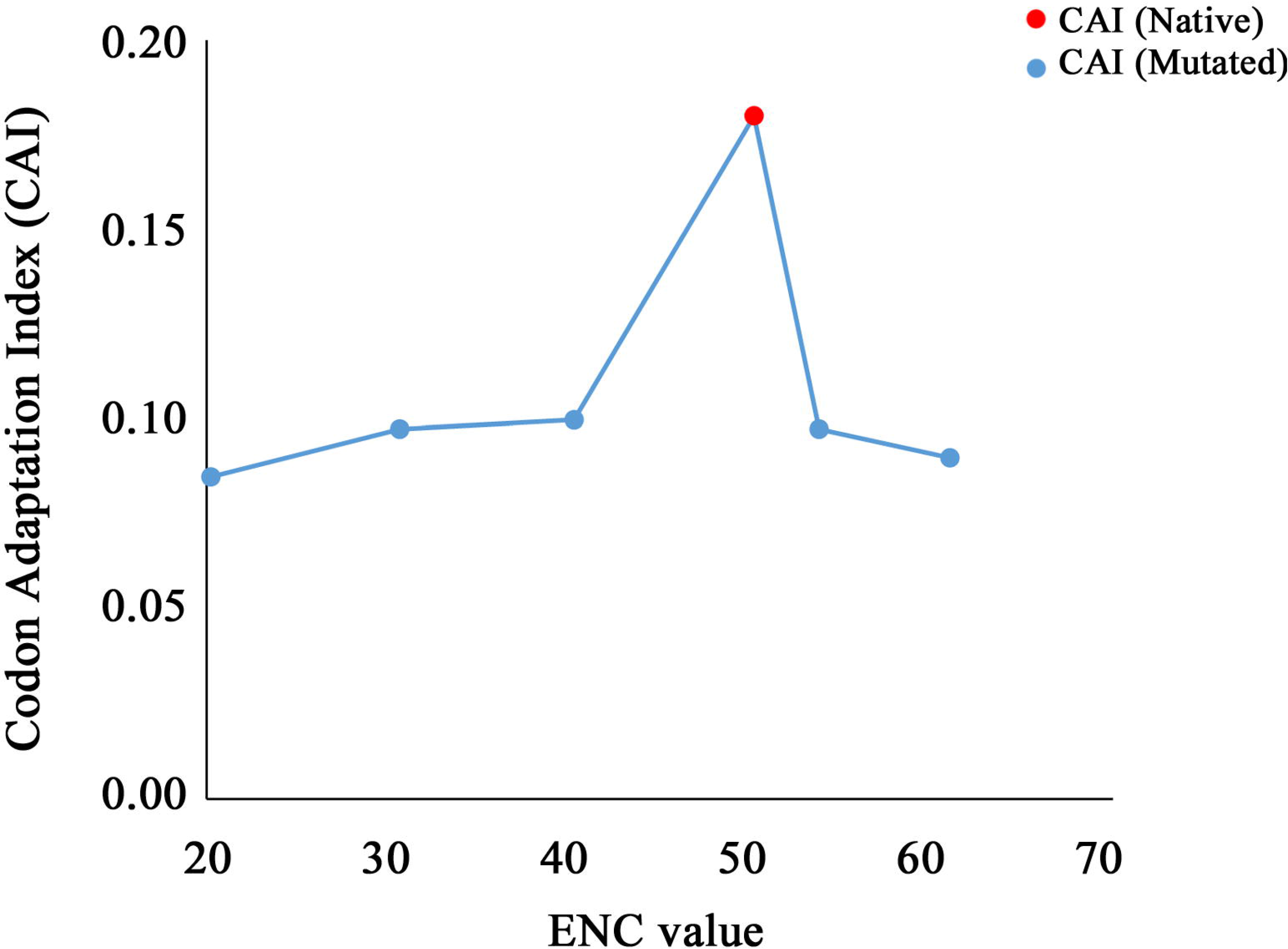
Average Codon adaptation indices (CAI) of *Saccharomyces cerevisiae* mRNAs in wild-type and the same generated computationally by different levels of codon compositions (mutants). The X-axis represents the ENC value for each group. The Y-axis represents the average CAI values for each group. The red dot represents the CAI and ENC in the wild type. Blue dots represent corresponding ENC and CAI values in mutants with different levels of codon composition.

### 3.4. Stability of native mRNA sequences and randomly shuffled sequences with fixed codon composition

The levels of gene expression, stability of mRNA secondary structure and codon usage bias have previously been shown to be strong predictors of translation levels and its associated attributes^12–14,16–18,21–23,37^. From our results it can be delineated that with increasing codon usage bias, mRNA secondary structure's stability is preferred and with stability on board elevated gene expression level is favoured. However, for a gene, the assimilated outcome of all these factors determines the gene expression levels and subsequently translation levels, which in this context of discussion favours increased levels^13^. However, we found that in such (both mRNA secondary structure stability and CUB is high) cases, gene expression does not increase, instead it becomes negatively correlated with mRNA stability under the fixed codon usage pattern (Supplementary table S1). As, for a gene, there are numerous theoretically possible combinations of mRNA sequences for the exact codon composition. If some codons are signatures for elevated gene expression, their presence will lead to higher expression of that gene (regardless of their positions), compared to genes that use such codons less often. In such cases, the gene expression is expected to remain unaltered, if codons present in the native structure are considered responsible for gene expression level. However, the stability of the mRNA is expected to vary, as with the change in the position of trinucleotides, structural alterations to this mRNA are expected to happen. We did a randomization experiment by interchanging the synonymous codons at different positions of the same mRNA of 5859 *Saccharomyces cerevisiae* genes. This lead to the generation of one hundred random sequences per mRNA that have similar ENC and CAI values. We calculated the stability of each mRNA for one hundred such randomly generated combinations, using both ΔG and folding score as measurements of mRNA stability (see Materials and Methods). We found that 5558 genes amongst 5859 showed the highest stability for native sequences i.e. 94.863% of total genes in a genome (Z score=97.128; p<0.0001). The result indicates the selection of stability for obtaining the apt mRNA abundance under the same ENC (Fig.2). This suggests that the native ENC values of the mRNAs helps to optimize the mRNA expression level, and among all possible codon composition of an mRNA, the structure with the highest stability is the native structure.

**Fig 2:**
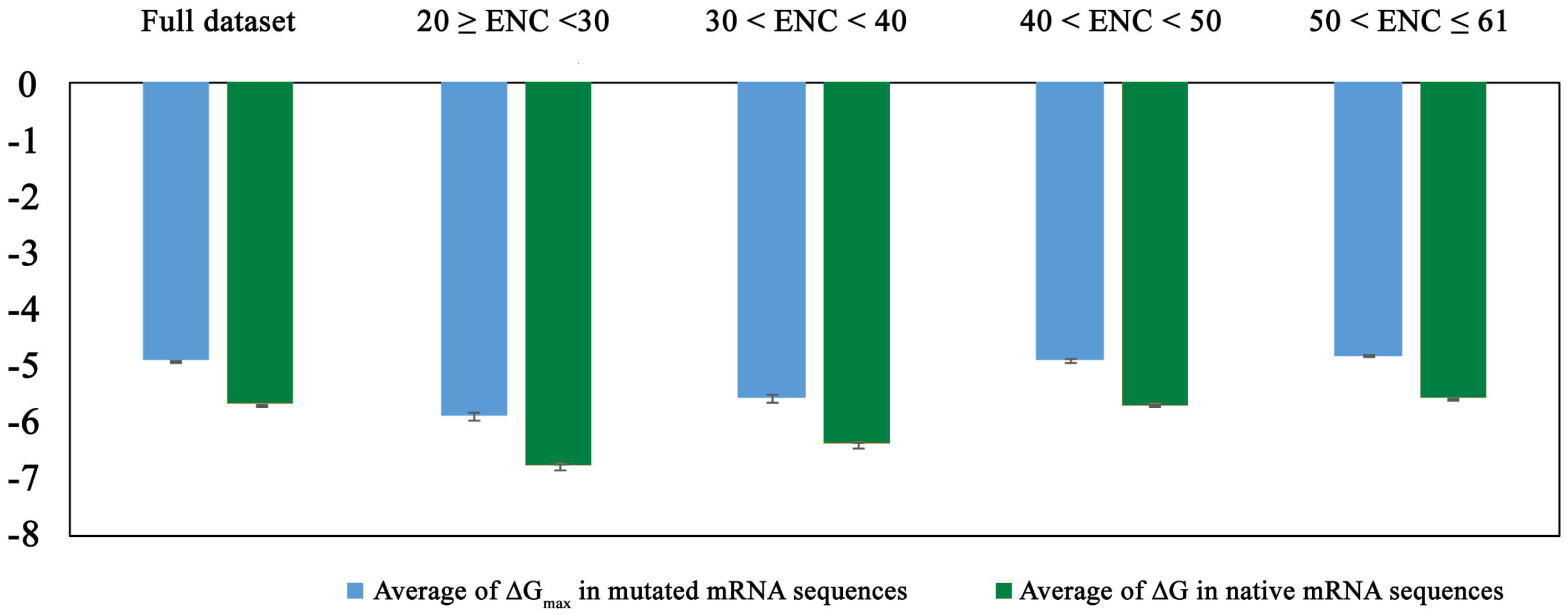
Comparison of mRNA stability in native and mutated mRNA sequences. In the whole dataset, as well as in each class based on native ENC values, the native sequence shows higher stability than the maximum stable mutated sequence.

Therefore, our study suggests that it is not necessarily correct that in a system if codon usage bias is raised even with high stability, the gene expression will be elevated. Rather, from the comparative analysis of mutational and native genome sequences, it becomes clear that there is demand for meticulous codon usage. If bias were the prime necessity then under the mutational study highly biased genome with high stability of the secondary structures would have elevated gene expression level. And as mRNA abundance holds a very strong positive relation with protein abundance in our datasets (ρ=0.614; P=1.0×10-6; n=4833) and (ρ=0.612; P=1×10-6; n=4374), it suggests that in this case (mutated genomes) there will be a downfall in the protein abundance. This result points in the direction that here is a choice of proper codon usage for appropriate mRNA stability, consequently contributing to mRNA expression and as well the modulation and regulation of translation articulates consonantly.

The transcriptional domain is essential because if stability is affected, the ribosomal binding will be affected and due to this, tRNA binding is also affected. Hence, translational rate and fidelity will be affected. So under the meticulous codon usage, the maximum expression is achieved along with maximum structural stability. This result also supports the positional bias of the codons under the light of structural stability, as codon positions were also found to be optimized for maximising structural stability. Together, these results indicate that the existing codon composition, rather than codon bias, is responsible for apt gene expression; and the naturally occurring mRNA structure in such a codon composition holds the key to maximum structural stability.

### 3.5. Evaluating mRNA abundance, mRNA stability, and codon usage bias

#### 3.5.1. Deciphering the relation between mRNA abundance, mRNA stability, and codon usage bias

From the correlation analysis, we have found that as the codon usage bias of the system increases, the stability of secondary structure enhances.

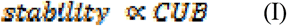

Similarly, the mRNA abundance and codon usage hold a strong positive correlation. It can be expressed as:

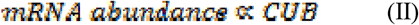

Combining equations (I) and (II) we find that

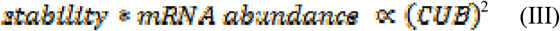

Now

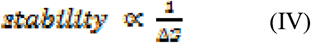

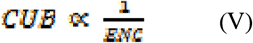

Using (III), (IV) and (V) we get

And

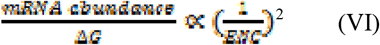

Multiplying both sides LHS and RHS by ΔG we get

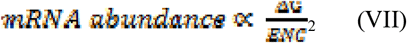

Irrespective of the bias (ENC) mRNA abundance will always be proportional to RHS.

If we assume a genetic system (genome), whose ENC doesn’t vary from one gene to another, then both stability and mRNA abundance will have no variation with ENC (I) and (II). So, using (I), (II), (VI) and (VII) it can be seen that under no variation of codon usage bias the gene expression level becomes inversely proportional to folding free energy. Hence, as the stability of the folded secondary structure increases the mRNA abundance will decrease. This is clear from the correlation analysis. But even under this condition, the mRNA abundance did not cease.

#### 3.5.2. Boolean expression combining ENC, mRNA stability, and mRNA abundance

Looking at the correlations we find that under different conditions of bias and hence stability as well, the mRNA abundance doesn't diminish to become nil, instead, lessens and behaves oppositely as compared to native system.

As gene expression level is a coordinated outcome of both codon usage bias and mRNA stability we tried expressing it as a Boolean expression.

With following considerations:

A: = CUB
B: = stability
C: = mRNA abundance
0: = low/absent
1: = High/Present

Making a truth table: see Table 4

**Table 4.** Truth Table showing the influence of mRNA stability and codon usage bias on mRNA expression level.

| C | A | B |  |
| --- | --- | --- | --- |
| 1 | 1 | 1 | When stability and bias both are present in high amount expression is there. |
| 1 | 1 | 0 | When stability is low and bias is high, expression happens |
| 1 | 0 | 1 | When bias is low and stability is high expression is present |
| 1 | 0 | 0 | When both bias stability are lowly present, expression is present |

As mRNA’s presence and its stability are co-existent so forming the equation with it.

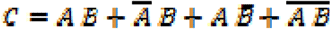

The realization of the logic gate is represented in **Fig.3**. The logical expression conveys that under all conditions we will have mRNA production (Table 4). Which means that codon usage pattern of any category (i.e. bias) and stability tends to produce mRNA, but its level is solely taken care by factor i.e. codon optimization as presented in the previous bias study. In a way it can be said that nature works for all codon composition but selects few for apt stability, abundance and translation regulation hence, codon optimization comes into the picture.

**Figure 3.**
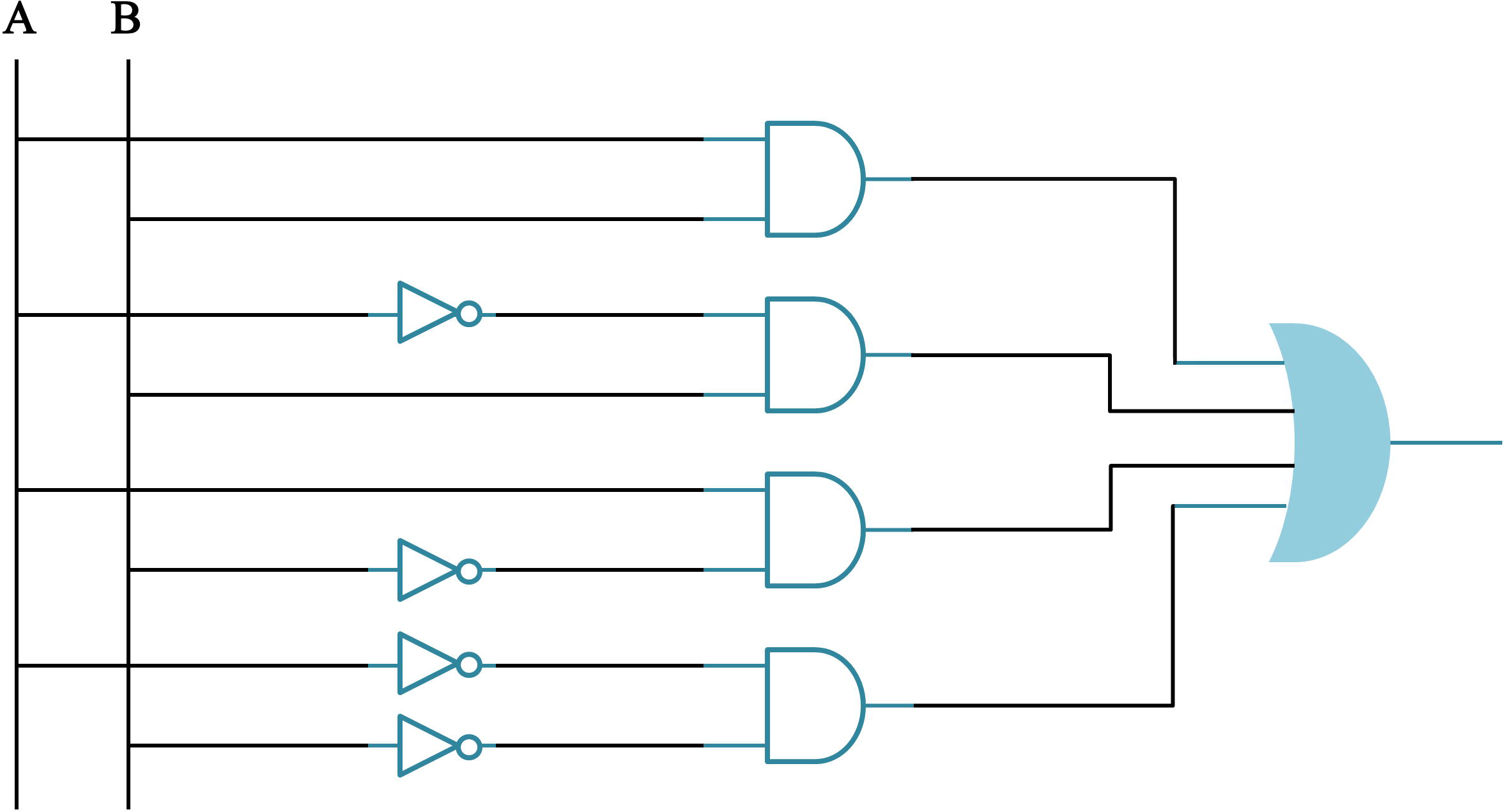
The logic gate realization of the Boolean expression stating the influence of mRNA stability and Codon usage bias on mRNA expression level.

## 4. Conclusion

Although biased usage of codons and high mRNA stability was long thought to be associated with elevated gene expression levels in the natural system, our results suggest that if any deviation is introduced to manipulate the naturally occurring codon composition, the gene expression always becomes reduced. The results show that even under the high stability and narrow selection of codon pool does not suffice for the appropriate selection for gene expression level, which is accounted from earlier studies. Additionally, our study also encompasses the positional bias of codons. We showed that even under native codon usage pattern there is selection for the most stable mRNA secondary structure. The results suggest a harmonious articulation of trio, i.e., codon composition, mRNA secondary structure stability, and mRNA abundance. Our study here confirms that the existent codon composition has not a mere influence of translational domain only, as transcription and translation are both articulated process. Rather, it indicates that both mRNA stability and meticulous codon usage fine tunes the apt gene expression level.

## Acknowledgements

The authors thank Mr. Sanjib Kumar Gupta and Dr. Jyotirmoy Das for their technical advice.

### Author contributions

MPV and TB conceived and performed the work. MPV and DA wrote the manuscript, DA, TB and TCG helped in planning and designing of the work and drafting the manuscript. All authors reviewed and approved the manuscript.

### Additional Information

### Competing financial interests

The authors declare no competing financial interests.

### Funding

This work was financially supported by Centre of Excellence in Bioinformatics, Bose Institute, Kolkata, India.

## Supplementary information

Supplementary file 1 containing Supplementary table S1.

